# Neural representation of bat predation risk and evasive flight in moths: a modelling approach

**DOI:** 10.1101/818609

**Authors:** Holger R. Goerlitz, Hannah M. ter Hofstede, Marc W. Holderied

## Abstract

Most animals are at risk from multiple predators and can vary anti-predator behaviour based on the level of threat posed by each predator. Animals use sensory systems to detect predator cues, but the relationship between the tuning of sensory systems and the sensory cues related to predator threat are not well-studied at the community level. Noctuid moths have ultrasound-sensitive ears to detect the echolocation calls of predatory bats. Here, combining empirical data and mathematical modelling, we show that moth hearing is adapted to provide information about the threat posed by different sympatric bat species. First, we found that multiple characteristics related to the threat posed by bats to moths correlate with bat echolocation call frequency. Second, the frequency tuning of the most sensitive auditory receptor in noctuid moth ears provides information allowing moths to escape detection by all sympatric bats with similar safety margin distances. Third, the least sensitive auditory receptor usually responds to bat echolocation calls at a similar distance across all moth species for a given bat species. If this neuron triggers last-ditch evasive flight, it suggests that there is an ideal reaction distance for each bat species, regardless of moth size. This study shows that even a very simple sensory system can adapt to deliver information suitable for triggering appropriate defensive reactions to each predator in a multiple predator community.

## 1. Introduction

Sensory system adaptations for detecting and responding to predator cues are well-known in animals, and the specific neural activity required to generate anti-predator behaviour has been documented for many species (e.g. arctiid moths: Ratcliffe et al. 2009; crickets: Nolen & Hoy 1984; locusts: Fotowat & Gabbiani 2007; cockroaches: Ritzmann 1980; crayfish: Edwards et al. 1999; fish: Eaton et al. 2001). Most animals co-occur with multiple predator species, each of which provides different cues and poses a different level of threat (Smolka et al. 2001; Falk et al. 2015). Animals often demonstrate anti-predator behaviours that are proportional to the risk posed by the detected predator (e.g. moths: Roeder 1974, Ratcliffe et al. 2011; crabs: Smolka et al. 2001; frogs: Fraker 2008; fish: Helfman 1989; birds: Templeton et al. 2005), but many aspects of predator threat cannot be directly encoded by sensory systems, such as the speed and manoeuvrability of a predator that is stationary at the time it is detected. Behavioural studies suggest that some animals might overcome this limitation by assessing a single detectable trait that is comparable across predators, such as size, to estimate predator threat (Templeton et al. 2005). Few studies, however, have assessed whether sensory systems are specifically tuned to cues that correlate with the predator threat posed by different predators in the community.

Eared moths and echolocating bats are ideal study animals for addressing this question. Echolocating bats are significant nocturnal predators of moths (reviewed in Fullard 1998). Bats produce ultrasonic echolocation calls for orientation and prey detection, and ultrasound-sensitive ears evolved in many moth families with the primary function of detecting bat echolocation calls (Fullard 1998; ter Hofstede & Ratcliffe 2016). With only 1-4 auditory receptor neurons depending on the moth family, these ears are the simplest ears in nature (Yack 2004). Moths in the family Noctuidae have two auditory receptor cells, called A1 and A2. The A2 cell is approximately 20 dB less sensitive than the A1 cell (**Fig. S1**). Noctuid moths also have a two-staged anti-bat response; they show directional flight away from quiet ultrasonic pulses, typical of a distant bat, and last-ditch flight manoeuvres in response to loud ultrasonic pulses, typical of a close bat (Roeder 1962, 1964; Agee 1969). Directional flight is likely triggered at intensities just above A1 threshold (Roeder 1964, 1967) and at much greater distances than those at which the bat can detect the moth’s echo (Roeder 1998; Surlykke et al. 1999; Goerlitz et al. 2010), meaning that moths can initiate directional flight to avoid being detected by the bat. Directional flight, however, should no longer be effective once the bat has detected the moth (Corcoran & Conner 2016) due to differences in flight speeds between bats (4-9 m/s: **Table S1**) and moths (0.5-6 m/s; Riley et al. 1992; Vickers & Baker 1997; Luo et al. 2002; Chapman et al. 2008; Corcoran and Conner 2016). Instead, moths require more drastic erratic flight or other secondary defence strategies (e.g., Ratcliffe & Fullard 2005, Corcoran & Conner 2012) to avoid being captured by the attacking bat. Due to the A2 cell’s higher thresholds, meaning it is only activated by high amplitude sounds, Roeder (1974) hypothesized that A2 cell activity might trigger these last-ditch flight maneuvers. This hypothesis, however, has never been empirically tested in noctuid moths.

The frequency tuning curves of moth A-cells vary depending on the sympatric bat community (reviewed in Fullard 1998; ter Hofstede et al. 2013), suggesting a strong functional link between frequency tuning of moth ears and the echolocation call frequency of sympatric bats. Although the A1 cell is more sensitive than the A2 cell, both cells have the same shaped tuning curve (**Fig. S1**), meaning that moths cannot discriminate between different bat species based on frequency (ter Hofstede et al. 2013). Larger moths have lower A1 cell thresholds (i.e. are more sensitive) than smaller moths, presumably to compensate for the greater distances at which bats can detect the louder echo reflected from their larger surface area (Surlykke et al. 1999; ter Hofstede et al. 2013). Despite differences in tuning and sensitivity between moth species, the general shape of the auditory tuning curve is similar across most moth species, with greatest sensitivity to lower ultrasonic frequencies and decreasing sensitivity at higher ultrasonic frequencies (**Fig. S1**; Fullard 1998; ter Hofstede et al. 2013; ter Hofstede & Ratcliffe 2016). The relatively consistent shape of the moth auditory tuning curve across species suggests that it is either constrained by morphological or physiological factors or that it is a functional adaptation, i.e. a matched filter (Wehner 1987; Römer 2016; von der Emde & Warrant 2016).

Here we use both empirical data and mathematical modelling to assess whether the shape of the moth auditory tuning curve is adapted such that moths detect different bat species at times when they pose a similar level of threat. First, we tested for relationships between bat call frequency, which varies across bat species in a community, and four variables that influence the detectability of calls by the moth and echoes by the bat (call intensity, duration and repetition rate) or the time required for the bat to intercept the moth (bat flight speed). Strong relationships between these variables would allow for the evolution of moth ear frequency tuning that reflects the threat posed by different bat species. Second, we incorporate data on bat echolocation call parameters, bat and moth hearing thresholds and estimated detection distances between bats and moths into multiple models of moth escape behaviour to develop testable hypotheses about the adaptive value of moth ear tuning for evading bat predators at a distance or at close range.

## 2. Relationships between bat call frequency and bat characteristics relevant to predator threat

We compiled data from the literature and our own measurements on echolocation call peak frequency, call duration, apparent call source level, call interval and flight speed for 14 European bat species (**Table S1**) that hunt flying moths (Barlow 1997; Vaughan 1997; Andersson et al. 1998; Bogdanowicz et al. 1999; Dietz et al. 2009). For our own measurements, we used acoustic flight path tracking to obtain these data for five European bat species (**Table S1**, for methods see Goerlitz et al. 2010). Thirteen of the bat species are in the family Vespertilionidae and produce relatively short duration calls at long intervals to separate call and echo in time, sometimes referred to as low duty-cycle echolocation (Jones & Teeling 2006; Fenton et al. 2012). One of the bat species (*Rhinolophus ferrumequinum*) is in the family Rhinolophidae, which uses a specialized form of echolocation, sometimes referred to as high duty-cycle echolocation, in which they produce long-duration calls and separate call and echo in frequency, allowing them to detect glints in the echoes caused by insects wing movements (Jones & Teeling 2006; Fenton et al. 2012). Species in this family, including *R. ferrumequinum*, often hunt insects close to vegetation (Jones & Rayner 1989; Dietz et al. 2009; Lazure & Fenton 2011).

For all bat species, we compiled data on call peak frequency (frequency with maximum energy), call duration (time from the start to the end of the call), apparent call source level (apparent emitted sound level at 10 cm from the bat’s mouth; i.e., the lower boundary of the source level because measurements also include off-axis calls), call interval (time from start of one call to start of the next call), and flight speed (the mean speed of multiple flight paths). We included these variables in our analyses due to their relevance to predator threat. Bats calling at lower frequency can detect prey, and can be detected by their prey, over greater distances because lower sound frequencies experience lower atmospheric attenuation than higher sound frequencies (Griffin 1971; Goerlitz 2018). Call duration was tested because moths are known to have lower hearing thresholds for longer duration sounds (Tougaard 1998), meaning that bats with longer calls would be detected by moths at greater distances than those with shorter calls. Likewise, bats that produce greater amplitude calls, measured as apparent call source level, can be detected by moths at greater distances than those with low amplitude calls. Long call intervals provide less information to the moth about predator positions than faster call rates. Finally, bats with faster flight speeds will close the distance between themselves and prey more quickly than bats with slower flight speeds. We used reduced major axis regression (Smith 2009; Trujillo-Ortiz & Hernandez-Walls 2010) to test for linear relationships between log10(call peak frequency) and the log10 of each of the four other bat characteristics. Major axis regression (also called geometric mean regression) is the appropriate method because both variables in the regression equation are random and subject to error.

Call duration, apparent call source level, call interval, and flight speed all had significant negative relationships with call peak frequency (**Fig. 1**, major axis regression, all p<0.05 after sequential Bonferroni correction). Bats calling at lower frequencies emit calls of higher amplitude and longer duration, call less often and fly faster than those calling at higher frequencies. These strong relationships between bat call frequency and bat characteristics relevant to predator threat suggests that the frequency tuning of moth auditory receptor cells might provide an adaptive filter for encoding predation threat across multiple sympatric bat species.

**Figure 1.**
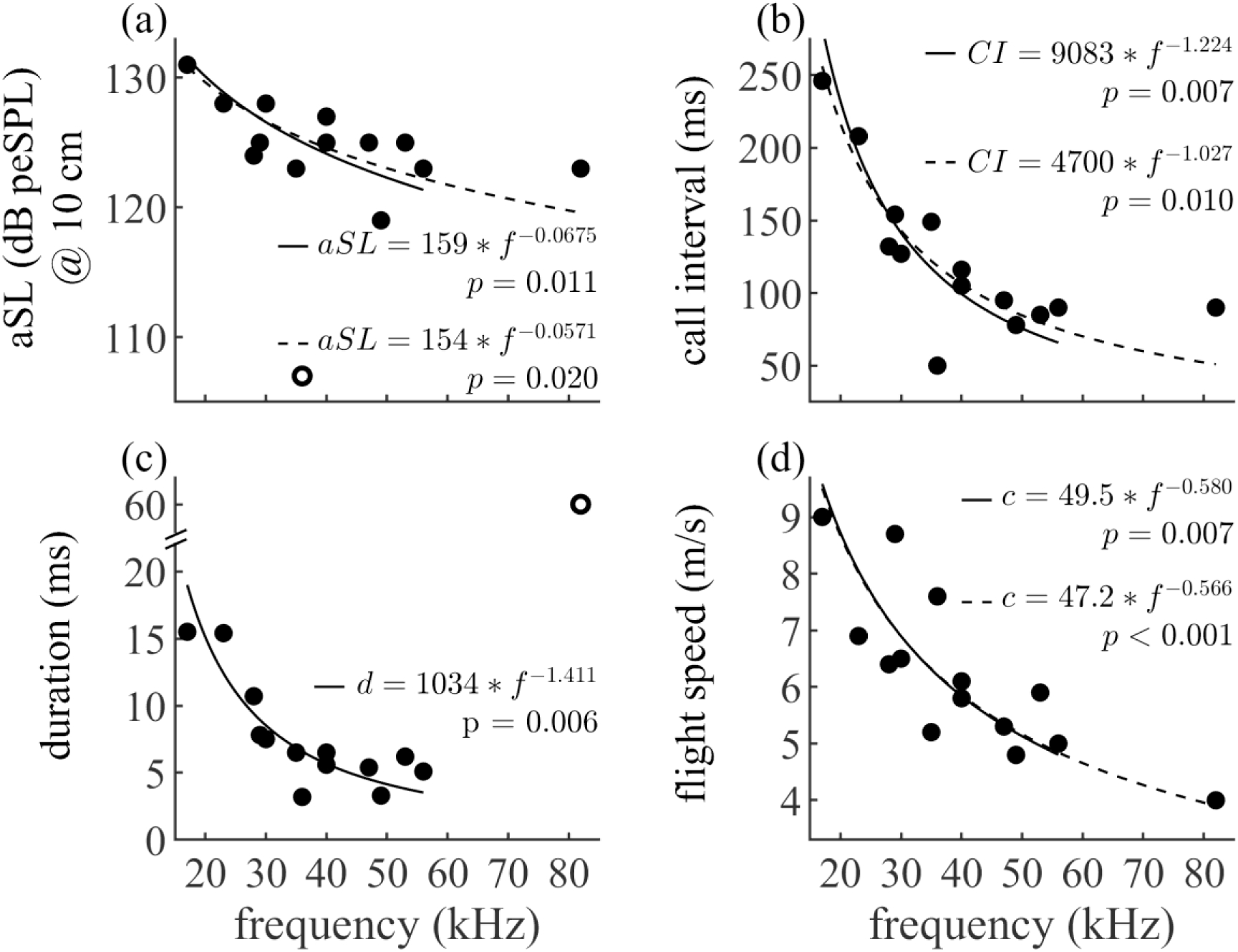
Bat echolocation call peak frequency predicts multiple characteristics of 14 European bat species: a) apparent source level (aSL), b) call interval, c) call duration, and d) flight speed. Solid and dashed lines show the relationship between bat characteristics and bat call frequency, including and excluding *R. ferrumequinum*, respectively. Note that we performed linear reduced major axis regressions of the logarithmized values, which we backtransformed to the here presented potential function of the non-logarithmized data. P-values report the results for testing the hypothesis that there is no relationship between the log-transformed data with sequential Bonferroni-correction for seven tests. We excluded the apparent source level of *B. barbastella* due to this species’ unusual low-amplitude stealth echolocation strategy (Goerlitz et al. 2010; Lewanzik & Goerlitz 2018) and the duration of *R. ferrumequinum* due to this species’ use of high duty-cycle echolocation (Schnitzler 1968) as outliers from all regression analyses (open circles in panels a and c). For data sources, see Table S1.

## 3. Calculating detection distances between bats and moths

To test whether the frequency tuning of the moth ear might function as a filter to match detection thresholds with the threat posed by different sympatric bat species, we calculated the maximum distances over which bats can detect moth echoes (bat detection distance, or bat-DD) and over which the A1 and A2 cells can be triggered by bat calls (A1-DD and A2-DD, **Fig. S2**). We calculated these values for all combinations of 14 European bat species (**Table S1**) and 12 European moth species (**Table S2**). Detection distances were calculated using the sonar equation (Møhl 1988), sound attenuation values based on mean weather conditions during our recordings (16°C, 75% relative humidity), estimated bat and measured moth hearing thresholds (see below), bat call source levels (**Table S1**) and moth target strength measurements (TS: echo level relative to the impinging sound level, **Table S2**; for details, see Goerlitz et al. 2010 supplementary methods).

Bat hearing thresholds have been estimated to be approximately between 0-20 dB SPL (Kick 1982, Neuweiler et al. 1984). Therefore, we set hearing threshold to 20 dB SPL for all bat species to account for intrinsic noise of the auditory system and (often even higher) behavioural reaction thresholds (Troest & Møhl 1986, Møhl 1988, Surlykke et al. 1999; Lewanzik & Goerlitz 2018). Moth hearing thresholds for bat calls were derived from published A1 and A2 thresholds for 20 ms pure tones (ter Hofstede et al. 2013; **Fig. 1**). First, we calculated the threshold at the bat species’ peak frequency by linear interpolation between the measured pure-tone thresholds at the frequencies below and above the bats’ peak frequency (for *R. ferrumequinum* calling at 82 kHz, we conservatively used the highest measured threshold at 80 kHz). Second, to account for differences in duration between pure tones and bat calls, we increased or reduced this threshold by 1.85 dB per halving or doubling of duration, respectively, to match the bat species’ call duration. This correction value is based on the relationship between sound duration and auditory threshold for two noctuid moth species (Tougaard 1998). These audiogram-derived A1 and A2 detection distances were additionally divided by 1.5 and 1.2, respectively. This is an accurate correction to obtain realistic estimates of maximum detection distances for bat calls in the field based on lab-measured audiograms (own unpublished data). Target strength was obtained from each moth’s surface-area as TS @ 10 cm = 13.5 dB * lg(SA) – 48.6 dB (Surlykke et al. 1999: Fig. 2a, mean of 30 and 100 kHz). Moth surface area was measured (ImageJ, National Institute of Health, U.S.A.) from digital photographs of moth specimens with wings completely spread (7±3 specimens per species (mean ± std), **Table S2**).

**Figure 2.**
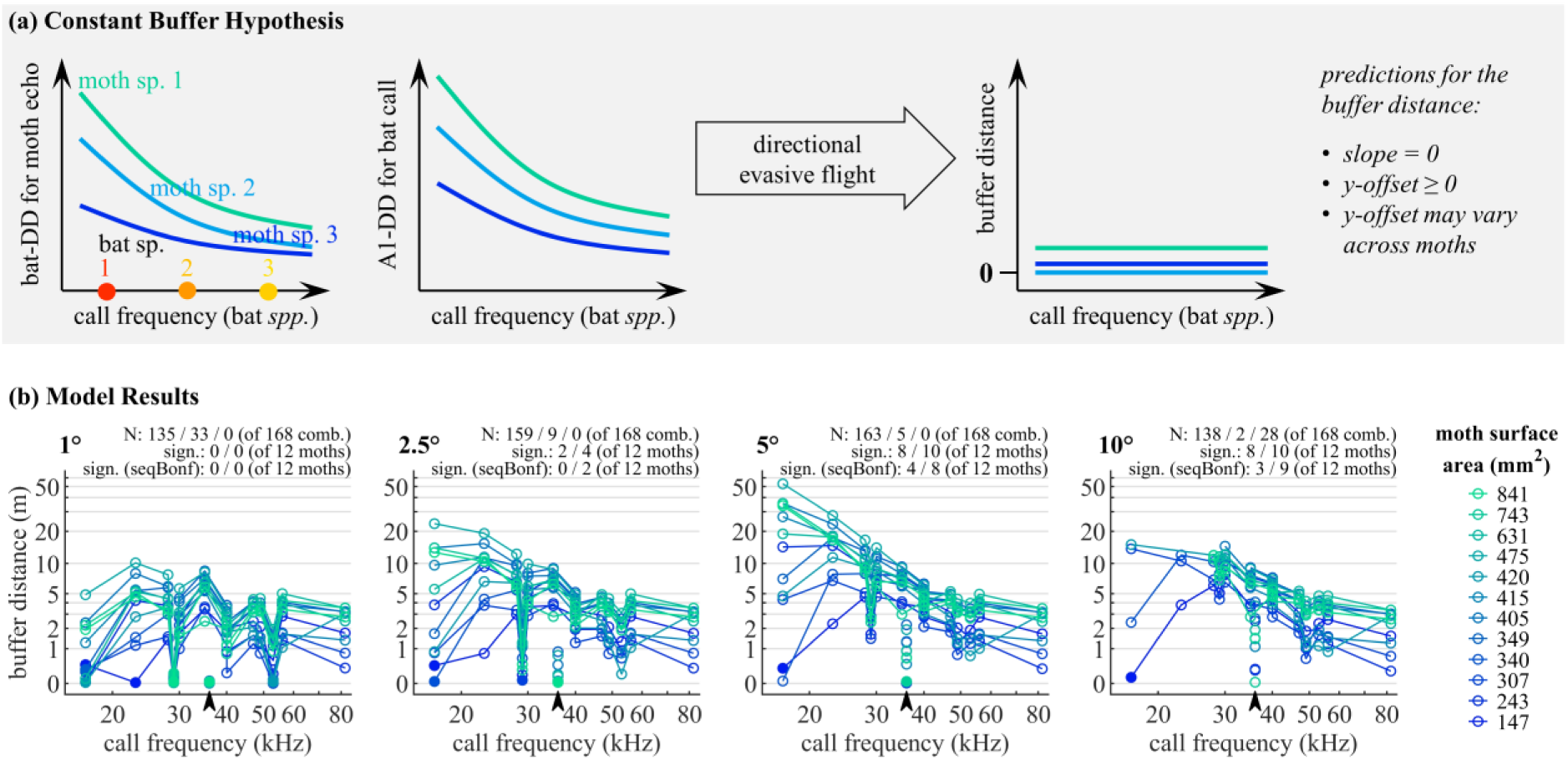
Predictions of the constant buffer hypothesis (a) and model results (b). **(a) Conceptual figure** showing the predicted results if the constant buffer hypothesis is correct and moths initiate directional evasive flight at the onset of A1 cell activity. The constant buffer hypothesis postulates that directional evasive flight allows moths to escape detection by a bat at a constant buffer distance before detection by the bat would have occurred, across all sympatric bat species for any given moth species. The curves in the first two panels symbolize the detection distances (DD) of moth echoes by bats (bat-DD), and of bat calls by the moths’ A1-cell (A1-DD), reflecting actual calculated values (see **Fig. S2**). The third panels shows the predicted constant buffer distance after directional evasive flight. **(b) Modelled A1-buffer distances** as a function of bat call frequency, and for four initial moth off-axis positions. Each coloured line represents one moth species. The initial off-axis position of the moth in relation to the bat flight and sonar beam direction is given at the top left of each panel. Open/closed symbols: moths that escaped detection and moths that were detected by the bat, respectively. Model parameters: 20° sonar beam width, 5 m/s moth flight speed. N: number of moth-bat-combinations (of 168) where the moth (i) escaped / (ii) was detected (incl. those where the moth’s initial position was within the bat’s search cone) / (iii) was outside the bat’s detection tunnel at the initial position and thus never detectable for the bat. Sign.: number of linear regressions (of 12 moth species) whose slope is significantly different from Zero, separately for regressions including / excluding *B. barbastellus*. Arrowhead: *B. barbastellus* (36 kHz), which was not connected by the coloured lines as it represents an outlier due to its exceptionally low amplitude calls.

## 4. Evading distant bats: modelling moth behaviour in response to A1 cell activity

### 4.1 The Constant Buffer Hypothesis

Noctuid moths show directional flight away from quiet pulses of ultrasound (Roeder 1962, 1964; Agee 1969). The A1 cell of the moth ear is more sensitive than the A2 cell (**Fig. S1**; Fullard 1998; ter Hofstede et al. 2013), and directional flight is likely triggered at intensities just above A1 threshold and below A2 threshold (Roeder 1964, 1967). Previous studies have shown that A1-DD is much greater than bat-DD (Roeder 1998; Surlykke et al. 1999; Goerlitz et al. 2010), meaning that moths can initiate directional flight before being detected by the bat. Bats with lower frequency echolocation calls can detects moths at greater distances than those with higher frequency echolocation calls, and most moths can detect bats with lower frequency echolocation calls at greater distances than bats with higher frequency echolocation calls (Surlykke et al. 1999). It is not known, however, how this correlation between detection distances relates to moth evasive behaviour.

The constant buffer hypothesis (**Fig. 2A**) suggests that the shape of the A1 tuning curve allows moths to detect bats with different echolocation call frequencies such that moths initiate directional flight in time to avoid detection by each bat species. Specifically, it predicts that if the shape of the moth auditory tuning curve is adaptive and the moth initiates directional flight away from the bat close to A1 threshold, then the moth will fly out of the path of an approaching bat with a similar buffer distance for each bat species.

### 4.2 Model design for the constant buffer hypothesis

To test the constant buffer hypothesis, we designed a geometric model that simulated linear bat search flight and directional evasive flight of moths upon detecting a bat call (cf., Corcoran & Conner 2016; Domenici 2002; Domenici et al. 2011; **Fig. 3**). Modelled bats flew along a straight path and emitted search calls of average duration, average call interval, average source level (**Table S1**), and half-amplitude beam widths of 15, 20, 25, 30 and 35° in the direction of flight (Ghose & Moss 2006) to cover documented variation (Jakobsen et al. 2012; Surlykke et al. 2009). We used the piston model (Surlykke et al. 2009) to calculate angle-dependent source levels (**Fig. 3c**). We used multiple angles of the moth’s initial position relative to the bat’s flight and sonar beam direction to determine how this off-axis angle would influence the moth’s escape probability (15 off-axis angles in total: 0.5, 1, 1.5, 2.5, 5, 7.5, 10, 15, 20, 25, 30, 40, 50, 70 and 90°). The initial distance between the modelled bat and moth was determined by the A1 detection distance (**Fig. 3a**), meaning that the received sound pressure levels of the bat calls were just at A1 threshold (Roeder 1964, 1967).

**Figure 3.**
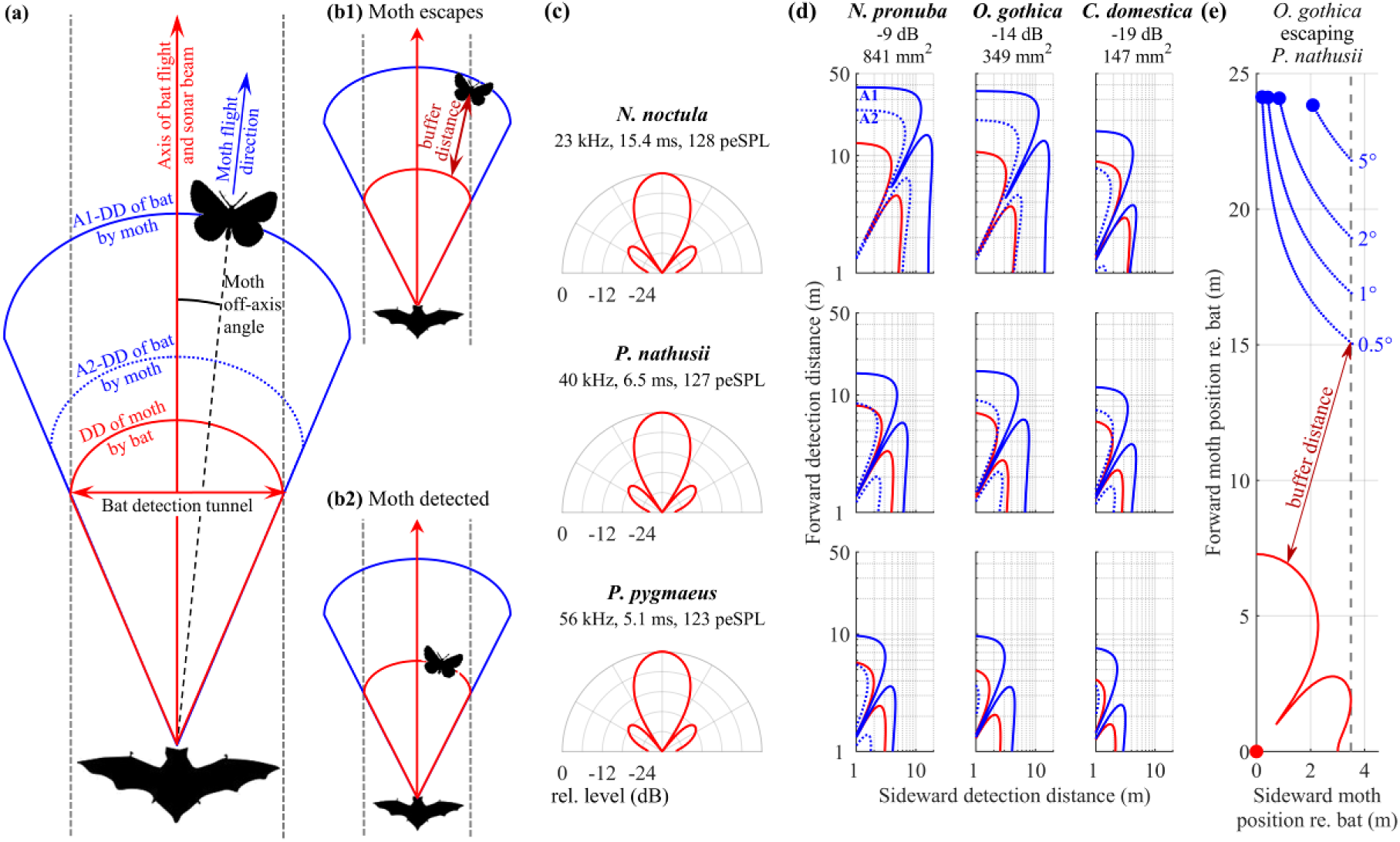
Diagrammatic representation of the model for directional evasive flight. **(a) Model features and terms.** Bats searched for prey along a linear trajectory, detecting any prey within their search cone as defined by their maximum detection distance (DD; red line). The maximum width of the bat’s search cone determines the bat detection tunnel (dashed grey lines), from which moths need to escape to avoid detection. The moths’ starting point was at the maximum detection distance of bats by their A1 cell (blue solid line) and at various off-axis angles relative to the bat’s flight trajectory. With each emitted call, bats flew along their linear search trajectory according to their flight speed, while moths flew directly away from the bat (negative phonotaxis) at various potential flight speeds. **(b) End points of the model.** The model run ended when the moth escaped by crossing the edge of the bat detection tunnel (b1), or when the moth was detected by crossing the bat’s maximum detection distance (b2). For escaped moths, the buffer distance is the shortest distance ever between any moth position and the front of the bat’s search cone (b1). **(c) Examples of sonar beam patterns** for three different bat species and 20° half-width beam angle, showing the call level emitted into all directions relative to the on-axis call level. While (a) and (b) show simplified sketches of the spatial search cone, (c) shows the realistic sonar beam pattern used in the model to calculate the realistic detection distances for the search cones in (d) and (e). Values below bat names are call peak frequency, duration and apparent source level (at 10 cm distance). **(d) Examples of maximum detection distances** of moths by bats (red) and bats by moths (blue, solid: A1, dotted: A2) for three different bat and moth species. Bat species are the same as in (c). Values below moth names are target strength (at 10 cm distance) and surface area. **(e) Example of a model run** showing moth positions (blue, each dot is a position per bat call) relative to bat position (red dot) for moth starting positions (large dot) at four different off-axis angles.

The model simulated the flight trajectory of the bat and moth with time steps equalling the call interval of the bat. After the emission of each call, bat and moth positions were altered by the distances they fly within one call interval. Bats flew with their average flight speed (**Table S1**) along their linear search path, starting with the first time step. Moths in the model initiated directional flight after the second call because moths only demonstrate sustained directional flight in response to multiple calls at repetition rates typical for bat echolocation (Roeder 1967). Moths moved directly away from the bat (i.e., along the bat-moth-direction = negative phonotaxis: Roeder 1962) at six different flight speeds (1, 2, 3, 4, 5 and 6 m/s), encompassing the range of measured noctuid moth flight speeds (Riley et al. 1992; Vickers & Baker 1997; Luo et al. 2002; Chapman et al. 2008; Corcoran and Conner 2016). After each call, we calculated the new bat-moth-distance, moth off-axis angle and respective detection distances for the new positions. The model run was stopped if (i) the moth escaped detection by the bat (i.e., it flew outside of the bat’s detection tunnel formed by the maximum lateral extent of the bat’s sonar beam and its forward movement; **Fig. 3b1**), or (ii) the moth was detected by the bat (bat-moth-distance was less than or equal to the maximum distance at which the bat can detect the moth; **Fig. 3b2**). If the moth escaped detection, we calculated the moth’s “buffer distance” as the shortest distance ever between the front of the bat’s echolocation search beam and the moth’s position during the entire modelled evasive flight (**Fig. 3b,e**). The buffer distance is a measure for the moth’s spatial safety margin, i.e., the shortest distance ever during evasive directional flight before the moth would have been detected by the bat. If the moth was detected by the bat, buffer distance was set to zero. If the moth’s initial position was already outside the detection tunnel, as was the case for many of the larger off-axis angles, then buffer distance was undefined.

We did not attempt to model the behaviour of bats and moths after the bat detected the moth for two reasons. First, bats have stereotyped echolocation while searching for prey, but change the structure of their calls and calling pattern once they have detected and pursue prey (Moss & Surlykke 2010), meaning that we could only effectively model the bats’ flight and calling behaviour prior to moth detection. Second, moths switch from directional evasive flight to more unpredictable last-ditch evasive flight while being approached by a bat. This switch might occur around the time when the bat detects the moth. The last-ditch behaviour varies widely, including loops, zig-zags and dives (Roeder 1962; Agee 1969), and varies between and sometimes within species (Hügel & Goerlitz 2019), making it currently impossible to model moth behaviour after the initiation of last-ditch behaviour. Therefore, we only modelled the interval during which we would expect to see moth directional flight away from the bat, i.e. after the moth detects the bat but before the bat detects and starts actively pursuing the moth.

For each moth species, our model resulted in one value of the buffer distance for each bat species. We evaluated the performance of the modelled evasive flight by calculating linear regressions between log10(buffer distance) and log10(bat echolocation call frequency). The log-transformation was used to linearize the data for statistical analyses. If the constant buffer hypothesis is correct, we predict that buffer distances within each moth species would be the same for all bat species regardless of echolocation call frequency, resulting in horizontal lines when buffer distance is plotted against frequency and a lack of significant regressions (**Fig. 2a**). We evaluated model performance by counting the number of linear regressions with slopes that are significantly different from zero (after sequential Bonferroni correction for twelve tests). Twelve insignificant regressions with slopes of zero would indicate strong support for the hypothesis, and 12 significant regressions would indicate rejection of the hypothesis.

### 4.3 Model results for the constant buffer hypothesis

We initially present results (**Fig. 2b**) for a typical sonar beam width of 20° (Surlykke et al. 2009; Jakobsen et al. 2012) and for a moth flight speed of 5 m/s, assuming that moths use a high flight speed during escape. Moths likely increase their flight speed when they hear bats, with evidence coming from observations of moths responding to bat attacks in free flight (Agee 1967, 1969; Corcoran & Conner 2016) and increased wingbeat frequency by tethered moths during their turning response to low amplitude ultrasound (Treat 1955; Roeder 1967). At small moth off-axis angles (up to 2.5°), buffer distance was constant across all bat species for most moth species (0 significant regression; the bat *B. barbastellus* was excluded because of its unusually low call source levels, **Fig. 2b**). With increasing moth off-axis angle, buffer distance increased for low-frequency bat species, resulting in fewer relationships that had constant buffer distances across bat species (only four and three moth species at 5° and 10°, respectively, excluding *B. barbastellus*). When the moth off-axis angle increased further, the initial position of a greater percentage of moths (10°: 17%, 15°: 48%, 20°: 74%) was already outside the detection tunnel, preventing the bat from ever detecting the moth.

Moth flight speed strongly influences the success of directional evasive flight (**Fig. 4a**, top to bottom). At slow moth flight speeds of 1-2 m/s, all moths flying in the centre of the bat detection tunnel (0-5° off-axis) are detected by all searching bats (blue line, **Fig. 4a**, top). At 10° off-axis, the proportion of bat-moth-combinations for which the moth escapes detection (green line) reaches almost 50% at 1 m/s and about 90% at 2 m/s moth flight speed. Increasing moth flight speed further allows more and more moths to escape from the centre of the bat detection tunnel without being detected by the bat. In contrast to moth flight speed, the bat’s sonar beam width has a very minor effect on the moths’ escape success (**Fig. S4**).

**Figure 4.**
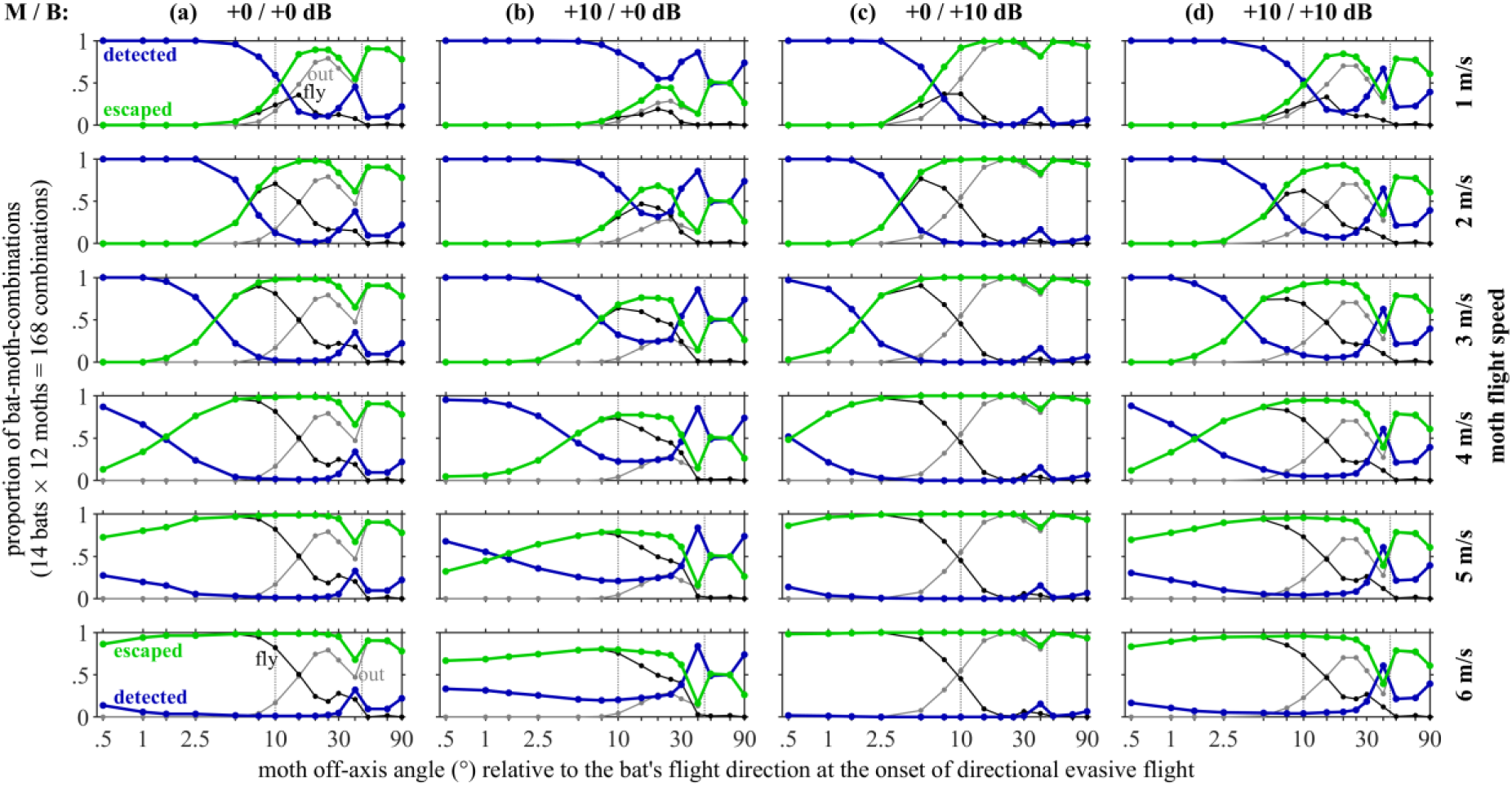
Effects of variation in behavioural threshold (a-d), moth flight speed (top to bottom) and moth off-axis angle (x-axes) on the success of directional evasive flight. Numbers above each column (+0 or +10 dB) are relative levels of reaction thresholds relative to moth A1 threshold (measured for each species) / bat hearing threshold (20 dB). Blue lines: proportion of moths that were detected by the bat; green lines: moths that escaped detection by the bat, either by flying out of the detection tunnel (black line, “fly”) or whose initial position was already outside of the detection tunnel (grey line, “out”). Dotted vertical lines are visual reference lines at 10° and 45° off-axis angle. Model parameters: 20° sonar beam width.

The behavioural reaction thresholds of bats and moths are generally not well-known, and might differ from neural detection thresholds. For moths, we used the neural detection thresholds of A1- and A2-cells as estimates for the onset of directional and erratic evasive flight, respectively. However, the exact mechanism linking neuronal activity to evasive flight is debated (Ratcliffe et al. 2009; Surlykke 1984; ter Hofstede & Ratcliffe 2016) and multiple acoustic call properties determine A-cell spiking and evasive flight (Roeder 1964; Waters 1996; Gordon & ter Hofstede 2018), which might only start at received levels of 10 dB and more above neuronal thresholds (summarized in Lewanzik & Goerlitz 2018). Likewise, bats’ reaction threshold can be 7 dB and more above our assumed threshold of 20 dB SPL (Lewanzik & Goerlitz 2018). We thus calculated all models again after increasing reaction thresholds by 10 dB for bats only, moths only and both bats and moths at the same time.

Increasing the reaction thresholds relative to the moth’s A1 detection thresholds and to 20 dB SPL bat threshold also influences the success of directional evasive flight. Increasing moth reaction thresholds by 10 dB generally lowers the proportion of moths that escape detection (**Fig. 4a,b)** from about 80% (135 of 168) to 45% (75 of 168, **Fig. S5a**, top two panels) at a moth flight of 5 m/s. Buffer distance of the escaped moths is somewhat lower, yet again constant across bat species (**Fig. S5a**), and the proportion of escaped moths increases with increasing moth flight speed as before with default thresholds (**Fig. 4b**). Likewise, a similar opposite trend exists when only increasing bat reaction threshold by 10 dB, allowing more moths to escape at larger buffer distances (**Fig. S5a**, third panel), lower speeds and lower off-axis angles (**Fig. 4c**). When increasing hearing thresholds of both moths and bats by 10 dB, the proportion of escaped moths, the buffer distance and the overall pattern is very similar to the default non-increased reaction thresholds (**Fig. 4d, S5a**). This suggests that the proportion of escaped and detected moths depends more on the relative difference in reaction threshold between bats and moths, and less on their absolute value, making our model results robust to variation in the exact reaction thresholds.

## 5. Evading close bats: modelling moth behaviour in response to A2 cell activity

### 5.1 The Matched Onset and Fixed Onset Hypotheses

To explain how the shape of the A2 cell tuning curve might be adaptive, we propose two mutually exclusive hypotheses. The matched onset hypothesis (**Fig. 5a**) postulates that last-ditch flight is initiated by the moth at the same distance at which the moth is detected by the bat (or possibly a constant spatial offset prior to that). In other words, using A2-DD as proxy for the initiation of last-ditch flight, this hypothesis postulates that the distance at which the moth A2 cell responds to bat calls (A2-DD) matches the maximum distance at which the bat can detect the moth (bat-DD). Thus, this hypothesis predicts 1) a slope of 1 for the relationship between A2-DD and bat-DD (overall, and also within each moth species and within each bat species); and 2) because bats detect larger moths over larger distances, i.e. bat-DD is positively correlated with moth size, A2-DD will also be positively correlated with moth size (**Fig. 5a**).

**Figure 5.**
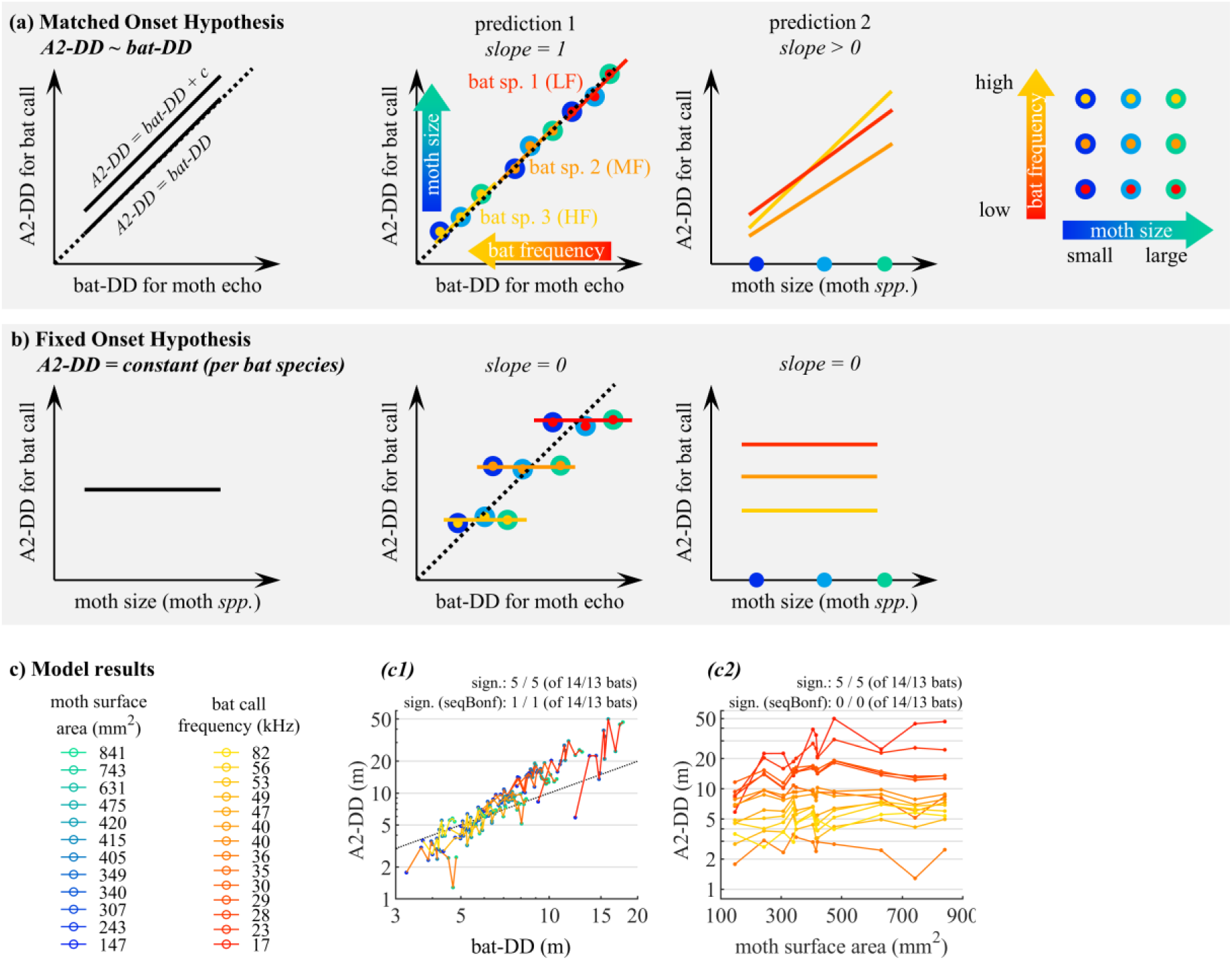
Predictions for the matched onset and fixed onset hypotheses (a,b) and model results (c). **(a,b) Conceptual figure** showing the predicted results if the matched onset hypothesis (a) or fixed onset hypothesis (b) is correct and moths initiate last-ditch evasive flight at the onset of A2 cell activity. **(a) The matched onset hypothesis** postulates that onset of erratic flight (approximated by A2-DD) is matched to the detection of the moth by the bat (bat-DD), resulting in a slope of 1 for the relationship between A2-DD and bat-DD, and a positive relationship between A2-DD and moth size. **(b) The fixed onset hypothesis** postulates that onset of erratic flight (approximated by A2-DD) is a fixed value for each bat species across all moth species, predicting no relationship between A2-DD and bat-DD (slope = 0), and no relationship between A2-DD and moth size (slope = 0). **(c) Model results:** the relationships between maximum detection distances of bat calls by the A2-cells (A2-DD) and of moth echoes by bats (bat-DD; c1), and between A2-DD and moth size (c2). Each colour represents one moth (blue to green) or bat (red to yellow) species support the fixed onset hypothesis. Sign.: number of linear regressions whose slope is significantly different from Zero, separately for regressions including / excluding *B. barbastellus*, for results before and after sequential Bonferroni correction. Arrowhead: *B. barbastellus* (36 kHz).

Alternatively, the fixed onset hypothesis (**Fig. 5b**) postulates that last-ditch flight might be most effective when initiated at a distance to impact that is specific to each bat species, not necessarily when the bat first detects the moth. This constant distance would be dependent on the flight speed and manoeuvrability of the attacking bat, but independent of moth characteristics. In other words, using A2-DD as proxy for the initiation of last-ditch flight, this hypothesis postulates that the distance at which the moth A2 cell responds to bat calls (A2-DD) is constant per bat species. Thus, this hypothesis predicts that 1) *within each bat species*, the slope of the relationship between A2-DD and bat-DD will be 0; and 2) *within each bat species*, A2-DD will be constant across moth species regardless of moth size (slope = 0), but this distance might vary between bat species (**Fig. 5b**). Both the matched onset and fixed onset hypotheses are based on the assumption that last-ditch flight is correlated with activity of the A2 cell, and this hypothesis has not been empirically tested in noctuid moths (ter Hofstede & Ratcliffe 2016). Our purpose is to determine how well the model results fit this assumption and to refine future hypotheses about the relationship between neural input and behavioural output in moths.

### 5.2 Model design for the matched onset and fixed onset hypotheses

To test the matched onset and fixed onset hypotheses, we tested for the existence of two relationships (i.e., having a slope significantly different from 0) within each bat species: 1) A2-DD vs bat-DD, and 2) A2-DD vs. moth surface area. We log10-transformed data before statistical analyses to achieve linearity. We assessed the support for each hypothesis based on the slopes and proportion of statistically significant regressions. First, the fixed onset hypothesis would be supported by significant positive relationships with a slope of 1 between A2-DD and bat-DD, and by significant positive relationships between A2-DD and moth size within bat species (**Fig. 5a**). In contrast, the fixed onset hypothesis would be supported by a lack of significant relationships and horizontal lines (slope of 0) between A2-DD and bat-DD and between A2-DD and moth size within bat species (**Fig. 5b**). We evaluated model performance by counting the number of linear regressions with slopes that are significantly different from zero (after sequential Bonferroni correction for 28 tests: 14 tests for A2-DD vs. bat-DD (one per bat species) and 14 tests for A2-DD vs. moth surface area (one per bat species)).

### 5.3 Model results for the matched onset and fixed onset hypotheses

The mutually exclusive matched detection and fixed onset hypotheses for the function of last-ditch evasive flight postulate that the A2-DDs are either matched to the bat’s detection distance of the moth, or alternatively are constant (fixed) for each bat species. For the test of prediction 1 (**Fig. 5c1**), approximately one third of the bat species (5 of 14 and 13) had significant relationships between A2 detection distance and bat detection distance. The five bat species with statistically significant slopes were the two bat species with the lowest call frequencies (*Nyctalus lasiopterus* and *N. noctula*) and three of the four bat species with the highest call frequencies (*Myotis* sp., *Pipistrellus pygmaeus* and *Rhinolophus ferrumequinum*). Only one of these relationships (*N. lasiopterus*) remained significant after sequential Bonferroni correction. Therefore, the A2 cells of all moth species start to fire at similar distances within almost all bat species, regardless of the bat’s detection distances for the moth, providing support for the fixed onset hypothesis and refuting the matched onset hypothesis.

For the test of prediction 2 (**Fig. 5c2**), approximately one third of the bat species (5 of 14 and 13) had significant relationships between A2 detection distance and moth size. The five bat species with statistically significant slopes were the two bat species with the lowest call frequencies (*Nyctalus lasiopterus* and *N. noctula*) and the three bat species with the highest call frequencies (*Miniopterus schreibersii, Pipistrellus pygmaeus* and *Rhinolophus ferrumequinum*). None of these relationships remained significant after sequential Bonferroni correction. Therefore, the A2 cells of all moth species start to fire at similar distances within those bat species, regardless of moth size, providing support for the fixed onset hypothesis and refuting the matched onset hypothesis.

Increasing the moths’ reaction threshold relative to the A2 cell detection threshold reduces the A2 detection distance for bat calls (**Fig. S5b2, second row**). Despite this, the number of non-significant relationships of A2-DD with moth size did not change, providing continued support for the fixed onset hypothesis. The number of significant relationships between A2-DD and bat-DD did not change when the moths’ reaction threshold was increased and increased from one to two significant relationships when the bat reaction threshold increased by 10 dB (**Fig. S5b1**). In summary, the fixed onset hypothesis is almost always supported for all tested moth and bat reaction thresholds.

## 6. Discussion

Our model results support the hypotheses that 1) echolocation call frequency of bats is correlated with variables that correspond with the predation threat they pose, 2) the shape of the A1 cell tuning curve is adapted to allow moths to avoid detection by bats with a similar spatial safety margin across bat species in their community, and 3) bats of the same species are detected by the moth A2 cell at a similar distance, regardless of moth species or size. In addition, the model suggests that the A1 cell is adapted for the worst-case scenario, allowing moths to avoid detection by bats when they are directly in the path of an oncoming bat, resulting in unnecessary defensive behaviour at greater angles away from a bat flying in a straight line. Together, these results suggest that the tuning of the moth ear is adapted to allow moths to respond to each bat species at an appropriate distance based on the frequency of its echolocation calls and suggest avenues for future empirical research and hypothesis testing.

### 6.1 Bat call frequency predicts threat-related bat characteristics

Larger bat species generally emit calls of lower frequency, greater call source level, longer duration and lower repetition rate than smaller bats (Barclay & Brigham 1991; Vaughan et al. 1997; Jones 1999; Holderied & von Helversen 2003). Our results demonstrate for the first time that many of these bat traits (call source level, call duration, call interval) as well as flight speed are tightly correlated with bat call peak frequency (cf. Vaughan et al. 1997; Waters et al. 1995 for call duration). These traits determine the maximum prey detection distance and the minimum time to prey interception, thus strongly influencing the risk that any given bat species poses for its prey. This correlation turns the bats’ (perceptible) call frequency into a proxy for its (imperceptible) predation threat. We suggest that the frequency-dependent tuning of moth auditory cells is a functional adaptation exploiting this correlation, allowing moths to respond to different sympatric bat species at appropriate distances.

### 6.2 Constant buffer hypothesis (onset of directional evasive flight)

Moths can detect bats at much greater distances than bats can detect moths (e.g. Holderied & von Helversen 2003; Goerlitz et al. 2010), giving moths the opportunity to initiate evasive flight before being detected. Our model of evasive directional flight based on measured moth auditory thresholds suggests that moth ears are adapted for the worst-case scenario, allowing moths to avoid detection by a bat even when they are in the centre of the bat’s flight path and echolocation beam. Remarkably, under these worst-case conditions, the buffer distance at which the modelled moths escape detection is relatively constant across bat species (0-10 m; **Fig. 2b**). This result suggests that the shape of the moth auditory tuning curve is adapted to allow moths to escape sympatric bats with a similar safety margin regardless of the frequency of a bat’s echolocation calls.

Since larger moths are detected by bats over longer ranges than small moths, Surlykke et al. (1999) already suggested that larger moths might need to detect bats at greater distances and require more time to escape the echolocation beam of bats than smaller moths. Our model supports this hypothesis, demonstrating that differences in detection range between large and small moths result in similar buffer distances of escape from the echolocation beam of an oncoming bat. Likewise, the different detection ranges for lower and higher frequency calls results in similar buffer distances of escape for low- and high-frequency bats.

Our model suggests that the buffer distances for escaping detection by a bat become more variable for larger off-axis angles of the moth relative to the bat’s flight and sonar beam direction. This large variation could be an accurate representation of buffer distances or it could be due to limitations of our model. If these values are accurate, moths detect lower frequency bats at much greater distances than necessary when they are off-axis from the oncoming bat. Unnecessary evasive flight reduces feeding and mating opportunities, but this might be an acceptable cost given the importance of avoiding predation. The simplicity of the moth ear likely constrains optimal adaptation to the full range of potential bat-moth orientations. On the other hand, our model limited the bat to fly in a straight line and to direct its echolocation calls straight forward, whereas foraging bats turn frequently and move their sonar beam to scan their surroundings (Surlykke & Moss 2000). The detection tunnel of a naturally scanning bat will be wider than modelled here, and the bat’s turns will increase the chances that a moth suddenly finds itself in the echolocation beam of a bat that is too close for effective escape by directional flight. The seemingly larger than necessary buffer distances might help moths to escape from turning and scanning bats. Lastly, the largest variation in buffer distances occurred at the lowest frequency (17 kHz), corresponding with the bat *Nyctalus lasiopterus*. We included *N. lasiopterus* to cover the full frequency range of European aerial hawking bat species, but it does not occur in the UK. Therefore, this bat species is not sympatric with the tested moth specimens in this study, which might result in relaxed selection at this frequency range for these moths (c.f. ter Hofstede et al. 2013). Another unknown is the directionality of hearing of bats, which is additive to the sonar beam directionality and will narrow the width of the bat’s detection tunnel. Hearing directionality is mostly unknown for bats, although for *Rhinolophus ferrumequinum* it is between 13-25° (Schnitzler & Grinnell 1977, Matsuta et al. 2013), i.e. in the range of the sonar beam directionality or even narrower. More data on hearing directionality, hearing direction, their variation due to ear movements (e.g., Griffin et al. 1962; Kugler et al. 2017, Lattenkamp et al. 2018, Yin & Mülller 2019), and on natural scanning behaviour will be required to estimate the average prey detection cone of free-flying bats.

Our model necessarily simplifies bat and moth behaviours and contains only a subset of the variables that could influence buffer distance. Future studies that collect data on the following variables would contribute to more accurate and precise models of bat-moth behaviour and help answer questions about the evolution of moth hearing for defence against bats. First, more detailed information on behavioural thresholds and how they relate to neural thresholds are needed. While correlative studies show that moths initiate directional flight at sound levels close to A1 cell threshold (Roeder 1962, 1967), direct measurements of A1 activity in behaving moths would help to confirm the exact relationship between neural activity and behaviour. Likewise, bat reaction thresholds are unknown and were set to 20 dB SPL, an estimate of hearing threshold for many bat species, although behavioural reaction thresholds can be higher (Lewanzik & Goerlitz 2018). Our analysis, however, shows that model results are rather robust against some variation in threshold and mostly depend on their relative values. Although our model did not include the moths’ reaction time (ranging between 45-250 ms from sound arrival to response onset; Roeder 1998), the model moths started their directional flight after the second bat call, reducing the influence of this variable on the results. Second, we assumed that moths would fly directly away from the sound source based on observational descriptions of moth behaviour (Roeder 1962), but three-dimensional flight path data are needed to confirm this. Third, incorporating species-specific data on moth flight speeds would improve the accuracy of the model, which we are currently missing for our species. Most studies that report flight speeds for moths focus on a few species of economic importance and measure flight in a pheromone plume. However, flight speed is often reduced within pheromone plumes and with increasing pheromone concentration (*Lymantria dispar*: Carde & Hagaman 1979, Charlton et al. 1993; *Heliothis virescens*: Vickers & Baker 1997; *Spodoptera littoralis*: Murlis & Bettany 1977). In addition, most studies investigated moth flight speed in wind tunnels, whereas moths increase flight speed with height above the ground (Keunen & Baker 1982) where the encounters between bats and moths would typically occur. Therefore, we only considered flight speeds that were measured in noctuid moths in the absence of pheromones and in free flight outside of a wind tunnel for the model. Lastly, flight speed is also likely positively correlated with moth size (*Lymantria dispar*: Kuenen & Carde (1993); butterflies: Dudley & Srygley (1994)), although this relationship is driven by weight and wing loading, not surface area. To overcome these limitations, we modelled all moth species at the same flight speed, but varied flight speed to determine its effect on the model results.

Despite these simplifications, our model predicts remarkably similar buffer distances across bat and moth species, suggesting that the tuning of the moth ear is a functional adaptation to the threat posed by bats with different call frequencies. Moths only escaped bats at higher flight speeds, providing specific predicted values for moth flight speeds while evading bats that can be tested in empirical studies. Future studies should consider collecting comparative data across a range of moth species of different sizes on 1) neural thresholds for turning responses, 2) 3D flight path tracking of moths evading distant moths (as measured by Corcoran and Conner 2016 for close range attacks), and 3) the relationship between moth size and flight speed in different contexts (both searching flight and directional evasive flight).

### 6.3 Matched onset and fixed onset Hypotheses (onset of last-ditch evasive flight)

Our model results provide support for the hypothesis that the A2 cell is adapted to fire at a fixed distance between a particular bat species and all sympatric moths regardless of moth species and size. If the A2 cell activity triggers last-ditch behaviour, this suggests that last-ditch flight might be most effective when initiated at a distance to impact that is specific to each bat species, not necessarily when the bat first detects the moth (as predicted by the matched onset hypothesis). In particular, it suggests that the ideal distance for moths to react to bats might be more influenced by aspects of bat flight speed and manoeuvrability than moth-specific parameters. Variation in the difference between A1 and A2 best thresholds across moth species of different sizes might contribute to this pattern. Although larger moths are more sensitive to ultrasound than smaller moths, the relationships between A1 threshold and moth surface area has a much steeper slope than that of A2 and moth surface area (ter Hofstede et al. 2013). Therefore, larger and smaller moths are more similar in A2 thresholds than A1 thresholds, contributing to this pattern of similar A2 detection distances seen across moths of different sizes in our model.

Despite almost complete support for the fixed onset over the matched onset hypothesis, there was sometimes still marked variation in A2-DD across moth species within a bat-species, particularly for low-frequency bats. Two factors contributed to this: bat-DD varied more with moth size for low-than high-frequency bats (**Fig. S6**), and A2-DD varied little with moth size for most bats, except for those with the lowest and highest call frequencies. Moth ears might be constrained to provide the most reliable representation of predator threat at intermediate frequencies, with bat species calling at extremely low or high frequencies having an advantage over moths, as proposed by the allotonic frequency hypothesis (Fullard 1998). Indeed, bats that call at extremely low or high frequencies are known to be moth specialists (*Euderma maculatum* 12 kHz: Fullard & Dawson 1997, Painter 2009; *Hipposideros ater* 160 kHz: Pavey & Burwell 1998).

Our model suggests two hypotheses for future empirical studies on the relationship between moth hearing and last-ditch anti-bat behaviour. Future studies should test whether A2 cell activity triggers last-ditch flight behaviour and whether this is true across all noctuid moth species or perhaps varies between species. Likewise, distances between bats and moths of known species and auditory sensitivity should also be measured with 3D tracking to confirm or refute that moths initiate last-ditch flight at a constant specific distance for each bat species.

### 6.4 Conclusions

To avoid responding to predators too late or too early, prey animals must correctly estimate the threat of multiple predators. Threat, however, depends on many predator-specific parameters and cannot always be directly perceived or assessed. In contrast, biophysical constraints in predators can result in correlations between particular predator cues that are detectable by prey, such as size or sound frequency, and the level of threat posed by a predator. Prey animals can benefit from sensory systems tuned to detectible cues in predators that provide information about potential predation risk across different predators. In echolocating bats, the sound frequency of their echolocation calls is a reliable and perceptible predictor of predation threat for eared insects. The frequency tuning of auditory receptor cells of eared moths appears to exploit this relationship, showing that even a very simple sensory system can adapt to deliver information suitable for triggering appropriate defensive reactions to each predator in a multiple predator community.

## Acknowledgements

Data collection was funded by the BBSRC (BB/f002386/1) to MWH. Model development and analysis was funded by the German Research Foundation (Emmy Noether, GO 2091/2-1 and GO 2091/2-2) to HRG. HtH was supported by funding from Dartmouth College. We are grateful to two anonymous reviewers for their valuable comments on an earlier version of this manuscript. We thank J. Memmott and G. Jones for assistance with collecting moths, Jim and Di McPetrie (Middlegrounds Farm Slapton), Slapton Ley Field Center, Mottisfont and Bristol City Council for site access and logistic support, D. Schröder and O. Nommensen for help with data analysis, L. Jakobsen for providing the Matlab code to estimate call beam shape, T. Hügel for discussion, and B. Hedwig, K. Kostarakos, T. Bayley, J. Memmott and G. Jones for constructive feedback on earlier versions of this manuscript.

## Data availability

DRYAD DOI: 10.5061/dryad.q5h132n

and temporary review link: https://datadryad.org/review?doi=doi:10.5061/dryad.q5h132n

## Author contributions

Devised the general research idea: HRG, HMtH, MWH

Collected and analysed moth neurophysiological data: HMtH

Collected and analysed bat data: HRG

Devised and developed the model: HRG, HMtH

Run and analysed the models: HRG

Wrote the manuscript: HRG, HMtH

Reviewed the final manuscript: HRG, HMtH, MWH

## Notes

**Declarations of interest:** none

#### Summary of Updates

A few minor changes througout the text: - corrections of a few typos - some subheadings are more precise - discussion on moth flight speed condensed - data points for Barbastella barbastellus in Fig. 2b not connected by lines anymore, as this species is an outlier due to its low-amplitude calls

