## Supplemental Tables and Figures for "Neural representation of bat predation risk and evasive flight in moths: a modelling approach"

### SUPPLEMENTARY TABLES

#### Table S1: Bat call and flight parameters used in the model

| | Call parameters | | | | |
|---|---|---|---|---|---|
| **Bat species** | **Peak frequency (kHz)** | **Duration (ms)** | **Apparent source level (dB peSPL) @ 10 cm** | **Call interval (ms)** | **Flight Speed (m/s)** |
| ***Nyctalus lasiopterus*** | 17 [2,4,5] | 15.5 [4,6] | 131 [2] | 246 [3,5] | 9.0 [3] |
| ***Nyctalus noctula*** | 23 [2,6,7,8,9] | 15.4 [6,7,8,9] | 128 [2] | 208 [3,8,9] | 6.9 [1,3] |
| ***Eptesicus nilssonii*** | 28 [2,6] | 10.7 [6] | 124 [2] | 132 [3] | 6.4 [3] |
| ***Nyctalus leisleri*** | 29 [1,2,6,7,8,9] | 7.8 [1,6,8,9] | 125 [1,2] | 154 [1,3,8,9] | 8.7 [1,3] |
| ***Eptesicus serotinus*** | 30 [2,6,7,8,9] | 7.5 [6,7,8,9] | 128 [2] | 127 [3,8,9] | 6.5 [3] |
| ***Hypsugo savii*** | 35 [2,6,8] | 6.5 [6,8,9] | 123 [2] | 149 [3,8] | 5.2 [3] |
| ***Barbastella barbastellus*** | 36 [1,6,7,8] | 3.2 [1,6,7,8] | 107 [10] | 50 [1,8] | 7.6 [1] |
| ***Pipistrellus nathusii*** | 40 [2,6,9] | 6.5 [6,9] | 127 [2] | 116 [3,9] | 5.8 [3] |
| ***Pipistrellus kuhlii*** | 40 [2,6,8] | 5.6 [6,8,9] | 125 [2] | 105 [3,8] | 6.1 [3] |
| ***Pipistrellus pipistrellus*** | 47 [1,2,6,7,8,9] | 5.4 [1,6,7,8] | 125 [1,2] | 95 [1,3,8,9] | 5.3 [1,3] |
| ***Myotis* spp.** | 49 [1,6,7,8,9] | 3.3 [1,6,7,8] | 119 [1] | 78 [1,8,9] | 4.8 [1] |
| ***Miniopterus schreibersii*** | 53 [2,6,8] | 6.2 [6] | 125 [2] | 85 [3,8] | 5.9 [3] |
| ***Pipistrellus pygmaeus*** | 56 [1,2,6,7,8,9] | 5.1 [1,6,7,8] | 123 [1,2] | 90 [1,3,8,9] | 5.0 [1,3] |
| ***Rhinolophus ferrumequinum*** | 82 [11,12,13,14] | 60 [11] | 123 [13] | 90 [11] | 4.0 [15] |

To account for variation in call and flight parameters, we used the mean of the values published in the following studies, as indicated by the small numbers:

[1] own unpublished data
[2] Holderied and von Helversen (2003)
[3] Holderied (2001)
[4] Estok and Siemers (2009)
[5] Ibanez et al. (2001)
[6] Obrist et al. (2004)
[7] Parsons and Jones (2000)
[8] Russo and Jones (2002)
[9] Vaughan et al. (1997)
[10] Lewanzik and Goerlitz (2018)
[11] Schnitzler (1968)
[12] Heller and von Helversen (1989)
[13] Schuchmann and Siemers (2010a)
[14] Schuchmann and Siemers (2010b)
[15] Dietz (2007)

For *N. noctula* and *B. barbastellus*, we used the mean of mean values of both call types (if available). For *Myotis* spp., due to the similarity in echolocation calls across species in this genus, we used the mean of *Myotis daubentonii* and *Myotis nattereri* from the literature.

**Table S2: Size (as surface area with wings fully spread) and calculated target strength of 12 UK moth species (Noctuidae) included in the model, ordered by size**

| | Surface area ($mm^2$) | | | Target Strength (dB) @ 10 cm |
|---|---|---|---|---|
| **Moth species** | **mean** | **sd** | **N** | |
| ***Cryphia domestica*** | 147 | 22 | 4 | -19.3 |
| ***Ochropleura plecta*** | 243 | 17 | 10 | -16.4 |
| ***Abrostola tripartita*** | 307 | 31 | 7 | -15 |
| ***Colocasia coryli*** | 340 | 48 | 5 | -14.4 |
| ***Orthosia gothica*** | 349 | 89 | 5 | -14.2 |
| ***Agrotis exclamationis*** | 405 | 29 | 10 | -13.4 |
| ***Xestia c-nigrum*** | 415 | 16 | 3 | -13.2 |
| ***Acronicta megacephala*** | 420 | | 1 | -13.2 |
| ***Autographa gamma*** | 475 | 44 | 7 | -12.4 |
| ***Apamea monoglypha*** | 631 | 55 | 10 | -10.8 |
| ***Amphipyra pyramidea*** | 743 | 83 | 7 | -9.8 |
| ***Noctua pronuba*** | 841 | 120 | 11 | -9.1 |

### SUPPLEMENTARY FIGURES

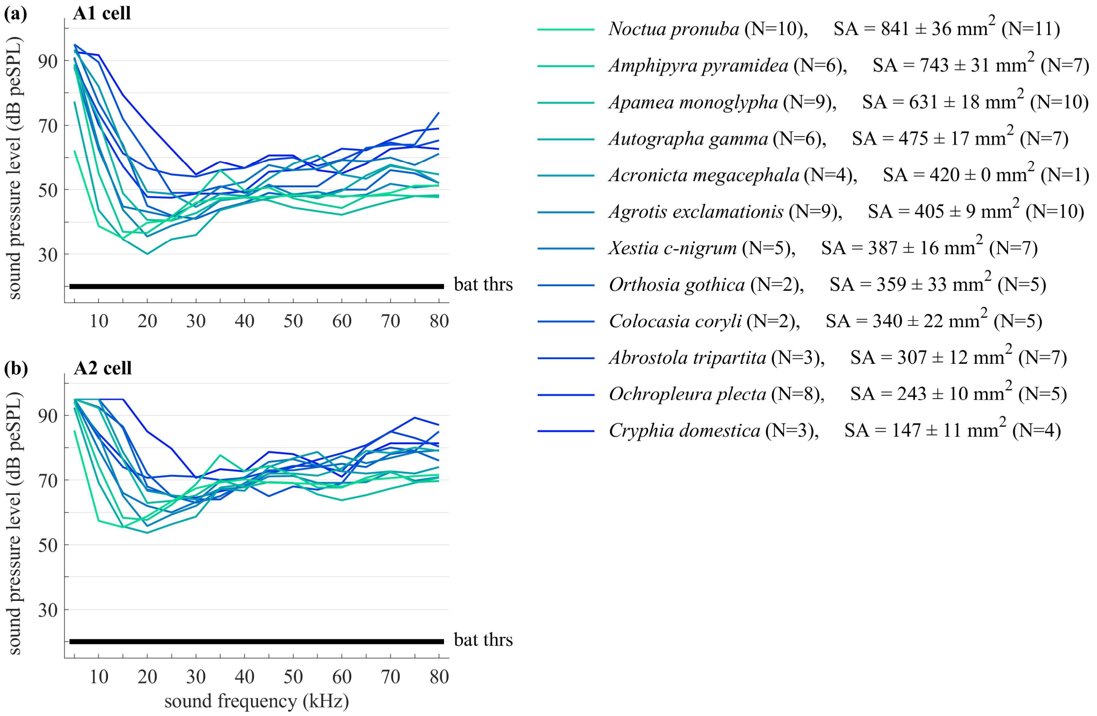


**Figure S1. Hearing thresholds of moths and bats.** Mean neural thresholds for the A1 (a) and A2 (b) auditory receptor cells in response to 20 ms pure tones for 12 moth species from the UK (data from ter Hofstede et al. 2013). Bat hearing threshold (black line) was set to 20 dB SPL for all bat species and frequencies. SA = surface area (mean ± s.e.m.); N after species name = number of moth individuals for neural data; N after SA data = number of moth individuals for SA data. Note that the mean audiograms shown here are 1.85 dB lower than shown in ter Hofstede et al. (2013). This difference is due to a correction applied by ter Hofstede et al. (2013) to recalculate the audiograms for shorter stimulus duration (10 ms instead of 20 ms).

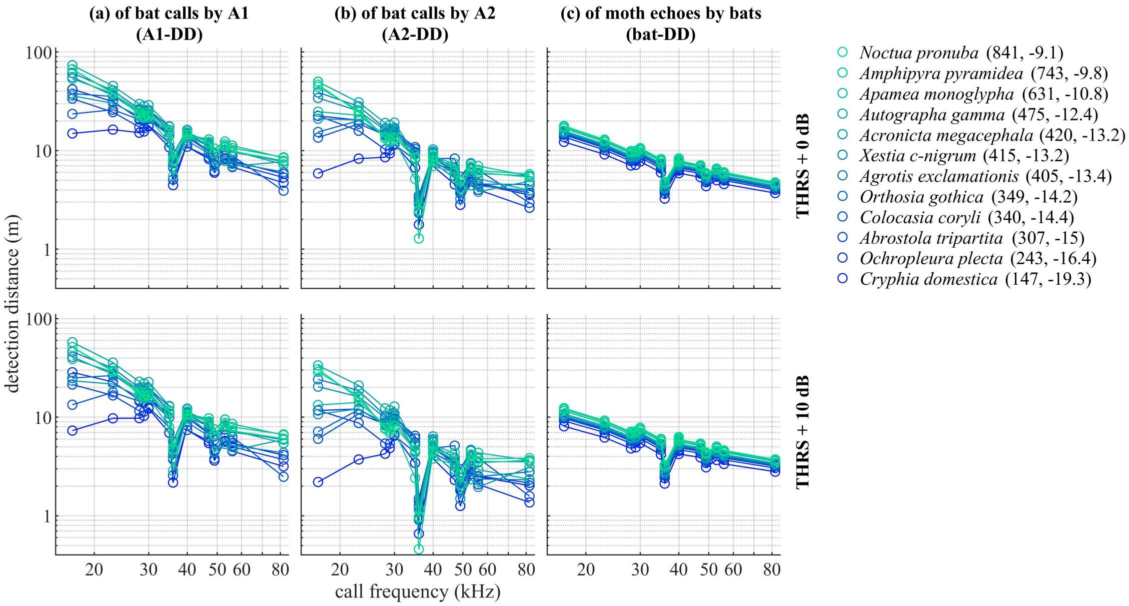


**Figure S2: Maximum calculated detection distances** of bat calls by the moths' A1 cell **(a)** and A2 cell **(b)**, and of moth echoes by bats **(c)**, for nominal hearing thresholds (top row) and for hearing thresholds increased by +10 dB (bottom row). Detection distances were calculated based on the bats' echolocation call parameters (**Table S1**) and hearing threshold of 20 dB SPL, the moth's surface areas (**Table S2**) and audiograms (**Fig. S1**), and for average weather conditions of 16°C and 75% relative humidity. Values in brackets after moth names state surface area (mm$^2$) and target strength (dB @ 10 cm distance to the moth; **Table S2**).

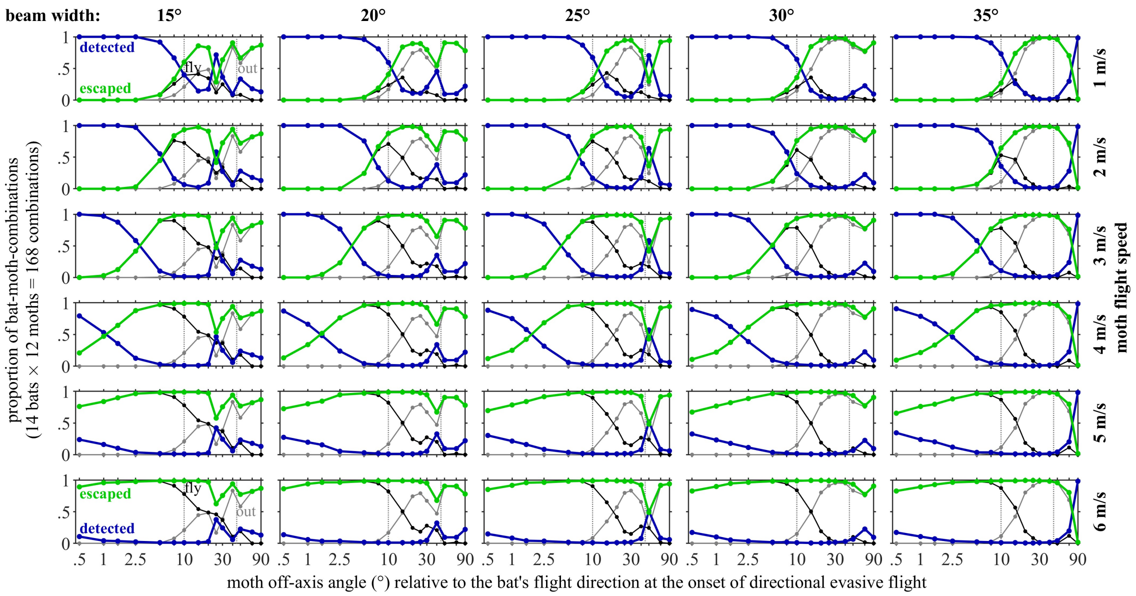


**Figure S4. Effect of half-amplitude sonar beam width (left to right), moth flight speed (top to bottom) and moth off-axis angle (x-axis) on the success of directional evasive flight.** Note how sonar beam width has only a minor influence on the success of directional evasive flight. In contrast, the influence of moth flight speed is much more pronounced.

Blue lines: proportion of moths that were detected by the bat; green lines: moths that escaped detection, either by flying out of the detection tunnel (black line, “fly”) or whose initial position was already outside of the detection tunnel (grey line. “out”).

Dotted vertical lines are visual reference lines at 10° and 45° off-axis angle. The model was calculated for default hearing thresholds (moth: A1 cell thresholds, bat: 20 dB SPL).

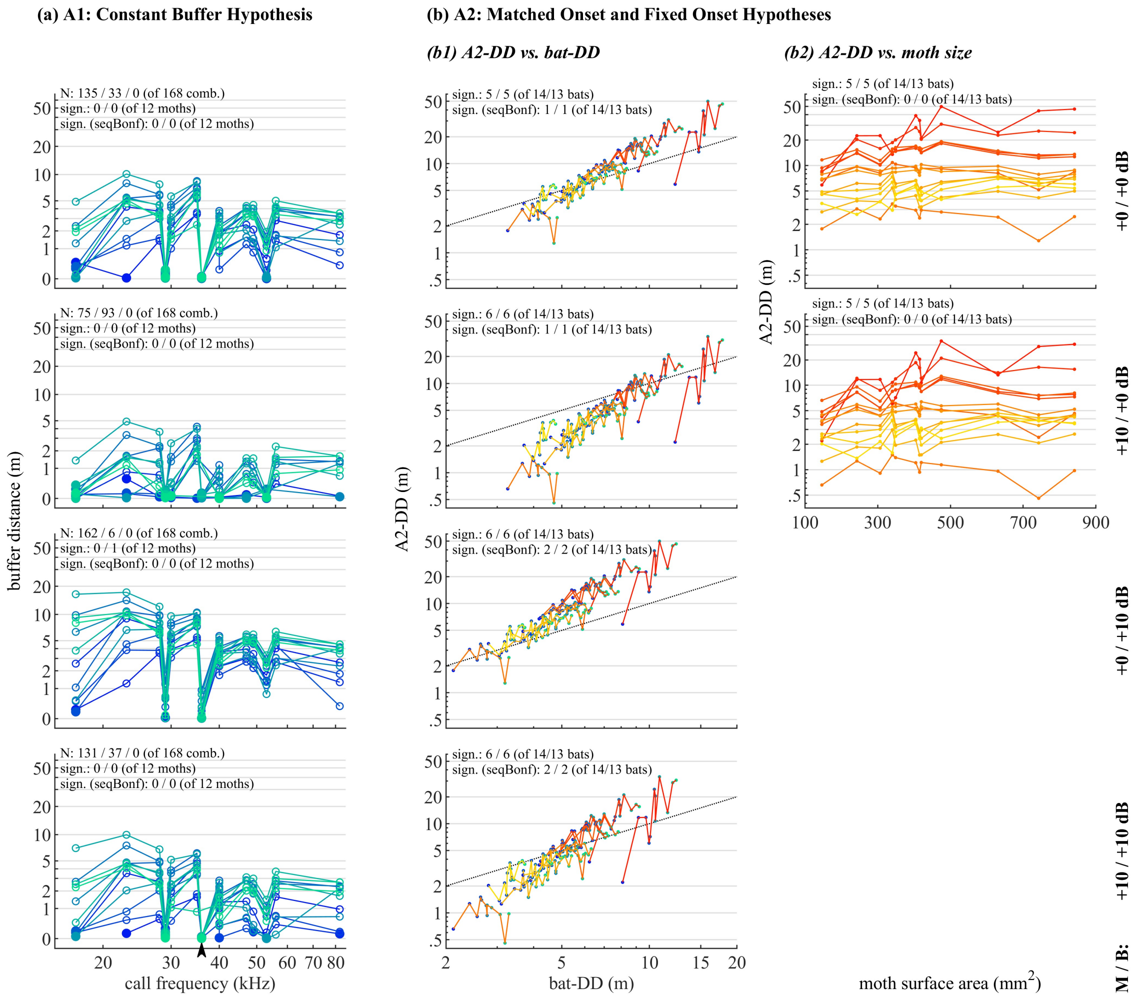


**Figure S5. Effect of variation in behavioural reaction thresholds (top to bottom) on the investigated variables of the Constant Buffer Hypothesis (a) and the Matched Onset and Fixed Onset Hypotheses (b).**

Numbers on the right of each row (+0 or +10 dB) are levels of the behavioural reaction thresholds relative to the moth A-cell threshold (measured for each species) and the bat hearing threshold (set to 20 dB SPL).

1° off-axis angle, 20° beam width and 5 m/s moth flight speed was used for (a). Note that A2-DD is independent of bat hearing threshold, thus results in (b2) are only shown once for default bat threshold (+0 dB).

N: number of moth-bat-combinations (of 168) where the moth (i) escaped / (ii) was detected (incl. those where the moth's initial position was within the bat's search cone) / (iii) was outside the bat's detection tunnel at the initial position and thus never detectable for the bat.

Sign.: number of linear regressions whose slope is significantly different from Zero, separately for regressions including / excluding *B. barbastellus*, for results before and after sequential Bonferroni correction. Arrowhead: *B. barbastellus* (36 kHz).

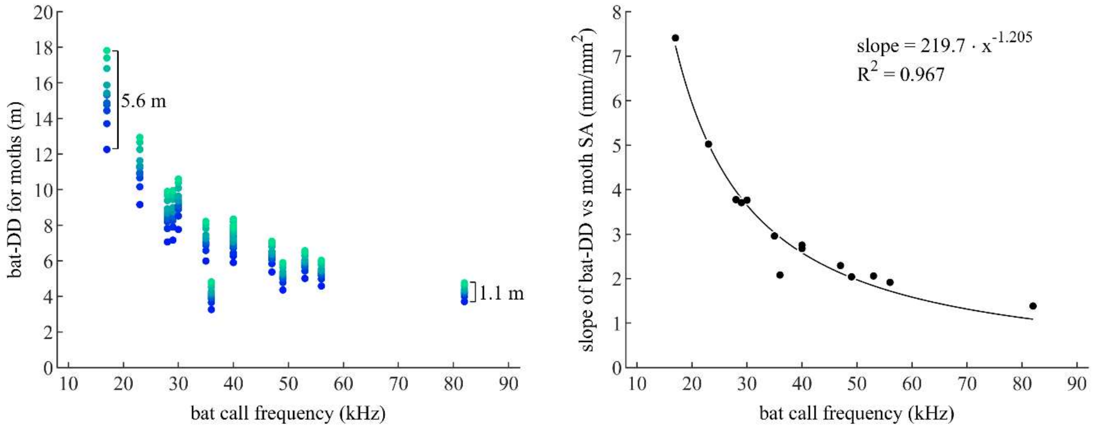


**Figure S6. Relationships between a) the distance at which bats can detect moths (bat-DD) and bat echolocation call frequency and b) the slope of the relationship between bat detection distance and moth surface area and bat call frequency.** Bat-DD varies by 5.6 m across moth species of different sizes for a low-frequency bat, while a high-frequency bat experiences a variation in detection distances for the same moths of only 1.1 m (a). This results in a larger slope of the relationship between bat-DD and moth size for lower frequency bats than for higher frequency bats (b).
